# De novo acetate production is coupled to central carbon metabolism in mammals

**DOI:** 10.1101/259523

**Authors:** Xiaojing Liu, Daniel E. Cooper, Ahmad A. Cluntun, Marc O. Warmoes, Steven Zhao, Michael A. Reid, Juan Liu, Kathryn E. Wellen, David G. Kirsch, Jason W. Locasale

## Abstract

In cases of nutritional excess, there is incomplete catabolism of a nutritional source and secretion of a waste product (overflow metabolism), such as the conversion of glucose to lactate (the Warburg Effect) in tumors. Here we report that excess glucose metabolism generates acetate, a key nutrient whose source has been unclear. Conversion of pyruvate, the product of glycolysis, to acetate occurs through two mechanisms: 1) coupling to reactive oxygen species (ROS), and 2) a neomorphic enzyme activity from keto acid dehydrogenases that enable it to function as a pyruvate decarboxylase. Furthermore, we demonstrate that glucose-derived acetate is sufficient to maintain acetyl-coenzyme A (Ac-CoA) pools and cell proliferation in certain limited metabolic environments such as during mitochondrial dysfunction or ATP citrate lyase (ACLY) deficiency. Thus, de novo acetate production is coupled to the activity of central carbon metabolism providing possible regulatory mechanisms and links to pathophysiology.

## Introduction

In conditions of hyperactive metabolism, excessive cellular nutrient uptake results in incomplete metabolism and excretion of waste products. For example, the carbon source for cellular energy-producing catabolic and biosynthetic anabolic processes is often incompletely catabolized and excreted into the extracellular space. During the Warburg effect, a phenotype characterized by increased glucose uptake, glycolysis rate (near 100 mM/h), is 10 to 100 times faster than the rate at which complete oxidation of glucose occurs in the mitochondria^1^. As a result, the excess carbon from glycolysis is secreted as lactate. Other examples are seen in the cells in diabetic tissues where excess fatty acid oxidation leads to incomplete lipid catabolism and excretion of acyl-carnitines and ketone bodies^2^. Numerous other examples are also apparent such as in the case of excess anabolic substrates released in nucleotide synthesis or the excretion of formic acid during excess one carbon metabolism^3^. In each case, the overflow metabolism results from imbalanced metabolic supply and demand and limits to enzyme activity.

The products of these overflow pathways provide a valuable resource during conditions of nutrient limitation. Ketone bodies become fuel sources during fasting conditions, while lactate and alanine are readily metabolized in local microenvironments that are deficient in glucose, amino acids, and oxygen^4–7^. These unconventional fuel sources satisfy metabolic demands during nutrient scarcity. Interestingly, acetate is also a nutrient that has been found to be a major carbon source for central carbon metabolism in nutrient limited conditions. Acetate metabolism provides a parallel pathway for acetyl-coA production separate from conversion of citrate to acetyl-CoA by ATP citrate lyase (ACLY) and thus acetate allows for protein acetylation and lipogenesis independent of citrate conversation to acetyl-CoA. This pathway is essential in nutrient deprived tumor microenvironments and other diverse contexts but the origin of acetate has been unclear^8–14^. It has been postulated that acetate may be synthesized de novo in cells^15–19^ but the pathways and quantitative reaction mechanisms through which this may occur are unknown. Given that alternative carbon sources often arise from overflow metabolism, we suspected that such a mechanism may allow for acetate generation. This hypothesis led us to conduct a re-evaluation of mammalian central carbon metabolism.

## Results

To monitor glycolysis in the setting of acetate metabolism, we incubated exponentially growing human colorectal HCT116 cells with uniformly labeled [^13^C_6_]-glucose, harvested metabolites over time, subjected the extracts to chemical derivatization with 2-hydrazinoquinoline (HQ) to capture acetate and other organic acids, and analyzed the products by liquid chromatography coupled to high resolution mass spectrometry (LC-HRMS) (Fig. 1a, methods). Monitoring the kinetics of [^13^C_3_]-pyruvate, [^13^C_2_]-acetate, and [^13^C_3_]-lactate, unexpectedly revealed that the rates of generation of pyruvate and lactate, the end products of glycolysis, were commensurate with the rate of acetate generation (Fig. 1b). Strikingly, the concentration of [^13^C_2_]-acetate was comparable to the concentration of [^13^C_3_]-pyruvate and on the order of the amount of [^13^C_3_]-lactate generated from [^13^C_6_]-glucose in these glycolytic cells. De novo acetate production from glycolysis was also observed in several other cell lines of diverse origins (Fig. 1c). Meanwhile, switching the ^13^C source from [^13^C_6_]-glucose to [^13^C_5_]-glutamine, which labels [^13^C_3_]-pyruvate and [^13^C_3_]-lactate to a far lesser extent, diminished the formation of [^13^C_2_]-acetate (Supplementary Figs. 1a and 1b). Together these findings suggest that acetate can be synthesized de novo from glucose in quantitative amounts and with kinetics comparable to pyruvate and lactate production from glycolysis.

**Figure 1.**
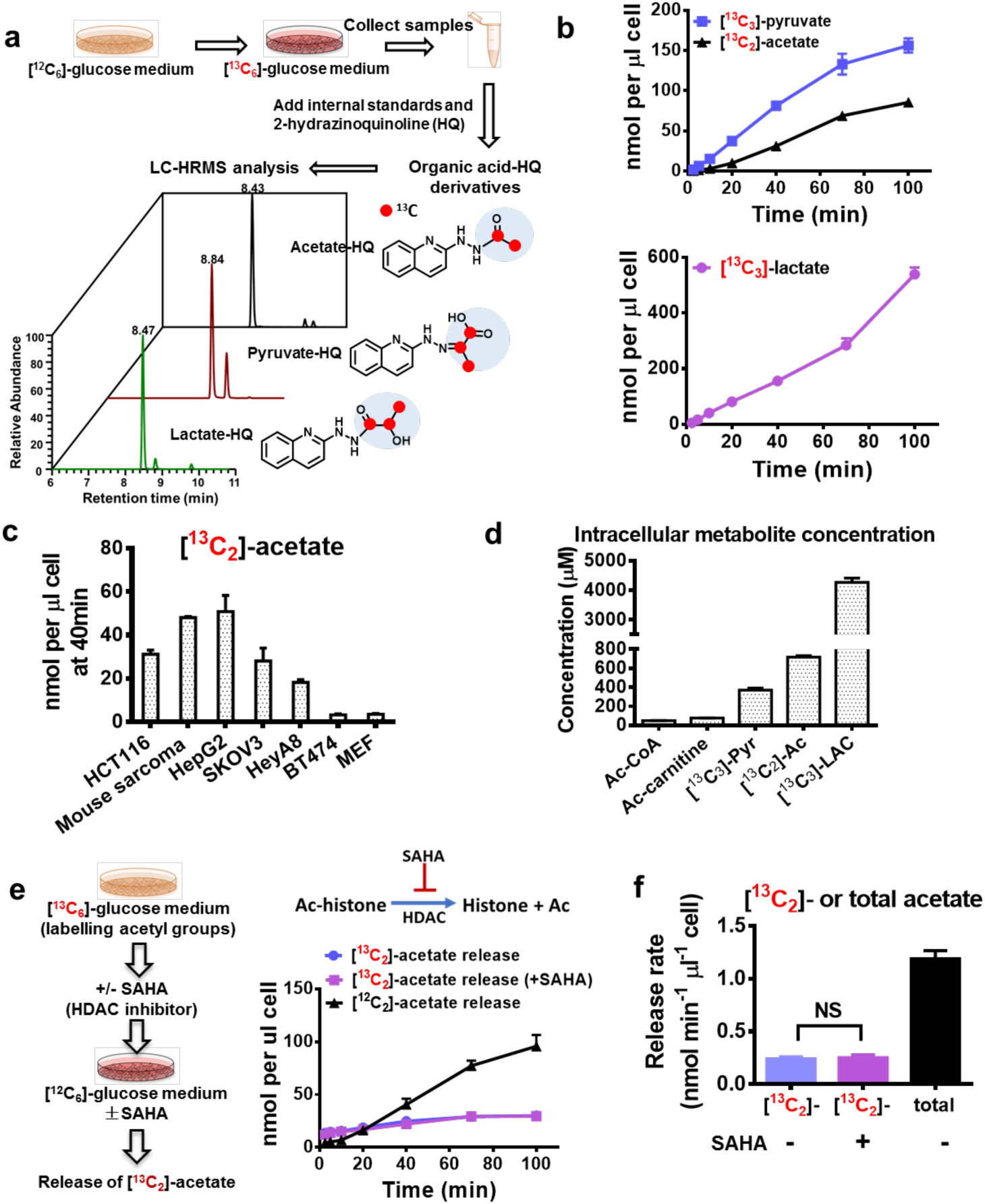
Acetate is synthesized from glucose independent of acetyl-CoA related reactions. **a,** Scheme of experimental setup for measuring glucose derived acetate, pyruvate and lactate. **b**, Medium concentration of [^13^C_3_]-pyruvate, [^13^C_2_]-acetate, and [^13^C_3_]-lactate secreted from HCT 116 cells cultured in RPMI 1640 medium containing [^13^C_6_]-glucose. Time 0 is when medium was switched to [^13^C_6_]-glucose medium. **c**, The amount of [^13^C_2_]-acetate released from various cultured cells at time 40 min after switching to [^13^C_6_]-glucose medium. **d**, Intracellular concentrations of metabolites in HCT116 cells. **e**, Release of acetate from HCT116 cells with 6 hours pretreatment of [^13^C_6_]-glucose in the presence or absence of histone deacetylation inhibitor, SAHA. **f**, The rate of [^13^C_2_]-acetate or total (^12^C and ^13^C) acetate release from HCT116 cells in the presence or absence of SAHA. The rate is obtained by calculating the slope of the acetate release curve (**e**). Ac: Acetate; Pyr: Pyruvate; Lac: Lactate; Ac-histone: acetylated histone. Error bars obtained from SD of n=3 independent measurements.

Acetate can be generated by the removal of acetyl groups from histones by histone deacetylases^20^ and by hydrolysis of Ac-CoA^21^. We thus measured the concentrations of the metabolites involved in these processes and observed orders of magnitude lower concentrations of Ac-CoA than glucose-derived pyruvate, lactate, and acetate (Fig. 1d), suggesting that de novo acetate production likely is uncoupled from these acetyl-CoA dependent reactions. Moreover, the acetate HQ derivative was not spontaneously formed by incubating Ac-CoA or acetyl (Ac)-carnitine with the HQ derivatization reagents indicating this potential artifact is sufficiently controlled (Supplementary Fig. 1c). To further evaluate the contribution of deacetylation reactions, including histone deacetylation, to acetate production, we cultured HCT116 cells in [^13^C_6_]-glucose containing media for 6 hours (hrs), which is thought to be sufficient to label histones with acetyl groups^22^, followed by incubation with the histone deacetylase inhibitor SAHA for an additional 1 hr (Fig. 1e). [^13^C_6_]-glucose medium was then replaced with ^12^C glucose medium, and the release of [^13^C_2_]-acetate would represent the contribution from acetyl groups to acetate production. Histone deacetylase inhibition didn’t alter the [^13^C_2_]-acetate release rate (Fig. 1e), and the measured [^13^C_2_]-acetate production rate from this assay was found to be nearly five-fold less than the total amount (^12^C and ^13^C acetate) of acetate (Fig. 1f), indicating that hydrolysis of the acetyl group from histones, known to be the most abundant source of deacetylation, is not a major source of acetate in these experiments. Together these experiments allow us to conclude that in these conditions, more than 80% of the acetate measured is rapidly generated de novo through glycolysis.

It is generally thought that Ac-CoA formation requires glucose to first enter the mitochondria, be exported as citrate and then metabolized to Ac-CoA (Supplementary Fig. 1d). We thus tested whether the pyruvate carrier inhibitor (UK5099)^23,24^ that perturbs entry of pyruvate into the mitochondria would affect acetate generation. As expected, the ^13^C labelled Ac-CoA, Ac-carnitine (reversibly converted from Ac-CoA) and citrate (the first intermediate in the TCA cycle) were decreased by UK5099 treatment (Supplementary Figs. 1e to 1g). Notably, no difference in the generation of [^13^C_2_]-acetate from [^13^C_6_]-glucose was observed (Supplementary Fig. 1h).

One considerable possibility of a substrate for acetate production is pyruvate, a keto acid, that contains an electrophilic moiety. Keto acids have been reported to be chemical scavengers in both bacterial culture and mammalian cells^25,26^. A plausible reaction pathway for the generation of acetate could involve the nucleophilic attack of pyruvate by the reactive oxygen species generated from hydrogen peroxide (H_2_O_2_) (Fig. 2a top), and the reaction would involve incorporation of the oxygen into H_2_O_2_ via production of superoxides and then into acetate. H_2_O_2_ thus obtains oxygen from molecular O_2_ and therefore, culturing cells in the presence of ^18^O_2_ and monitoring incorporation of ^18^O into acetate would enable quantitation of the endogenous contribution of ROS to acetate production. In this experimental design, other potential acetate production routes (deacetylation or aldehyde oxidation) would involve transferring the oxygen in a water molecule (which is negligible in this setup) to acetate and thus these two possibilities could be resolved with this experiment (Fig. 2a bottom). Therefore, we cultured cells in the presence of 20% O_2_ (^18^O_2_ and ^16^O_2_) and 80% N_2_ (Fig. 2b). We then employed LC-HRMS to enable spectral resolution of [^18^O_1_]-acetate from other isotopically labeled species. Analysis of the incorporation of ^18^O into tryptophan-related metabolites that are produced through oxygenation and stable labeling after 48 hrs demonstrated that 45% of the total O_2_ is ^18^O labeled in this setup (Fig. 2c). We then identified (Fig. 2d) and quantified (Fig. 2e) [^18^O_1_]-acetate providing a direct observation *in vivo* of formation of [^18^O_2_]-H_2_O_2_ and oxidative decarboxylation of pyruvate by H_2_O_2_. To further test this mechanism, an inhibitor of superoxide dismutase (SOD), tetrathiomolybdate (TTM), which converts superoxides to H_2_O_2_ (Fig. 2f), was used to decrease endogenous H_2_O_2_ levels. Indeed, the presence of TTM decreased ^18^O incorporation into H_2_O_2_-coupled reactions, including methionine sulfoxide (Fig. 2g) and acetate (Figs. 2 h and 2i). These findings together demonstrate that endogenous H_2_O_2_ contributes to acetate formation from pyruvate in cellular conditions.

**Figure 2.**
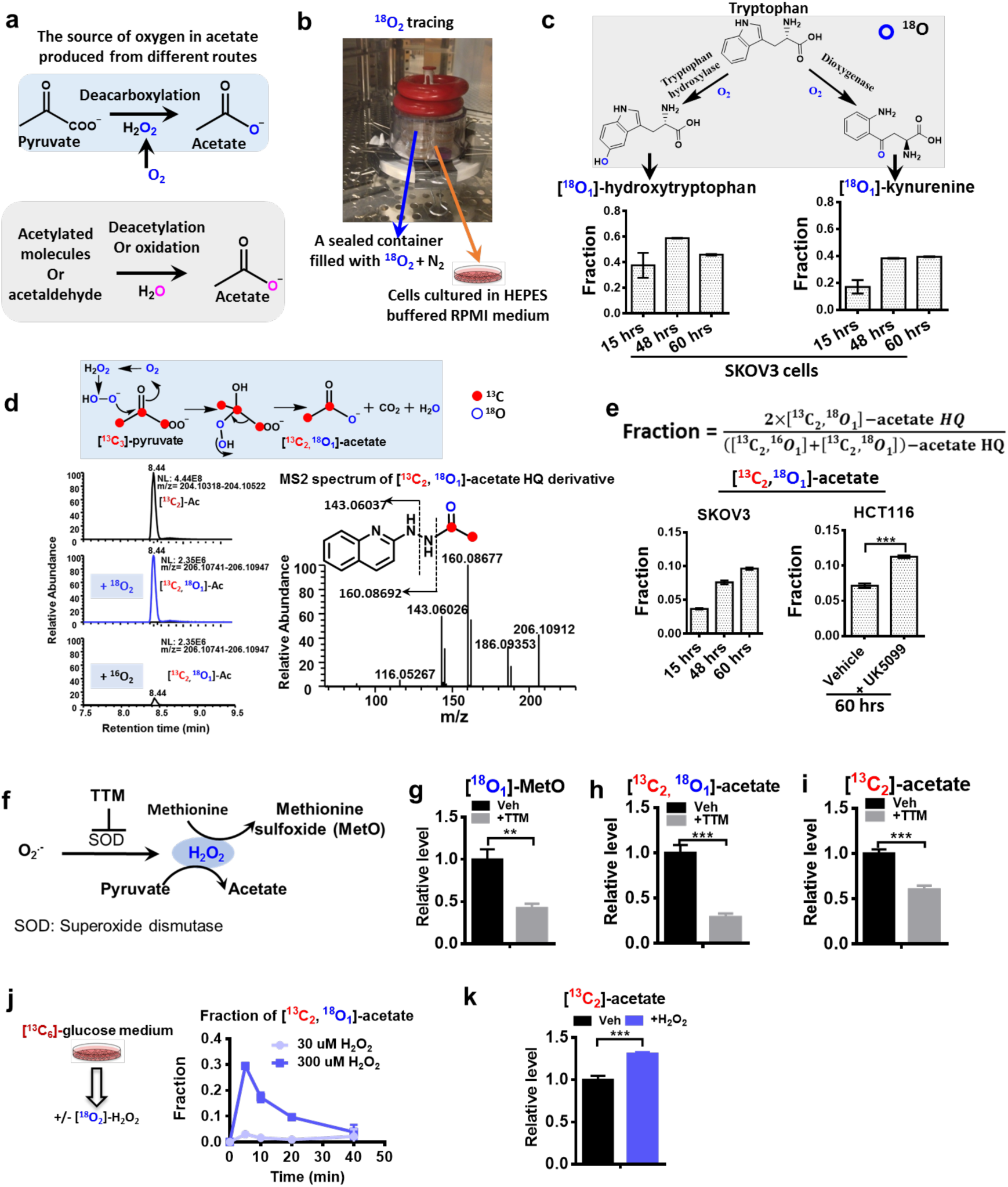
ROS catalyzes the oxidative decarboxylation of pyruvate to acetate in cancer cells. **a**, Model of the potential source of oxygen in acetate produced from different routes. **b**, Experimental setup of ^18^O_2_ tracing assay. **c**, Incorporation of ^18^O into metabolites in tryptophan metabolism as a marker to assess the ^18^O_2_ tracing assay. **d**, Scheme of ^18^O and ^13^C incorporation into acetate produced from H_2_O_2_ mediated pyruvate decarboxylation, and representative LC chromatography and MS2 spectrum of [^13^C_2_, ^18^O_1_]-acetate derivative. Blue open circle and red solid circle denote ^18^O and ^13^C, respectively. For simplicity, the O in carbonyl group of acetate HQ derivative is blue color highlighted, but due to the resonance structures in carboxyl group, the ^18^O could be either in the carbonyl and hydroxyl group. **e**, Fraction of [^13^C_2_, ^18^O_1_]-acetate out of glucose derived acetate pool. **f**, Scheme of endogenous H_2_O_2_ generated from superoxide and superoxide dismutase (SOD) and H_2_O_2_ mediated methionine oxidation and pyruvate decarboxylation. TTM, Ammonium tetrathiomolybdate, a SOD inhibitor. The effect of SOD inhibitor TTM on the relative level of [^18^O_1_]-MetO **(g)**, [^13^C_2_, ^18^O_1_]-acetate **(h)** in SKOV3 cells cultured in [^13^C_6_]-glucose and ^18^O_2_ for 48 hrs. **i**, Secretion of [^13^C_2_]-acetate from HCT116 cells in the presence or absence of TTM. **j**, Release of [^13^C_2_, ^18^O_1_]-acetate from HCT116 cells after addition of exogenous [^18^O_2_]-H_2_O_2_. **k**, Relative abundance of [^13^C_2_]-acetate and [^18^O_1_, ^13^C_2_]-acetate in either cell free medium or spent medium collected at 10 min after addition of 0 or increasing concentrations of [^18^O_2_]-H_2_O_2_ to HCT116 cells which were pre-incubated in [^13^C_6_]-glucose medium for 1 hour. Error bars obtained from SD of n=3 independent measurements.

To further investigate the properties of this reaction, we performed *in vitro* assays by incubating pyruvate with hydrogen peroxide at 37 °C and found the major product to be acetate as measured by ^1^H Nuclear Magnetic Resonance Spectroscopy (Supplementary Fig. 2a). Thus, in the presence of hydrogen peroxide, pyruvate is converted to acetate non-enzymatically with kinetics commensurate with values needed for the reaction to occur at appreciable amounts in cells (Supplementary Fig. 2b). The reaction followed second order kinetics (k = 0.19 +/- 0.05 mM^-1^ min^-1^) and could be accelerated with catalysts present in high concentrations in cells such as Cu^2+^, as Cu^2+^ stabilizes the intermediate of pyruvate decarboxylation (Supplementary Figs. 2b and c).

As a further evaluation, we tested whether exogenous H_2_O_2_ affects acetate production by adding [^18^O_2_]-H_2_O_2_ to cultured HCT116 cells (Fig. 2j). Titrating concentrations of [^18^O_2_]-H_2_O_2_ revealed a dose-dependent increase in acetate levels and the acetate detected had ^18^O incorporation as measured by LC-HRMS (Fig. 2j). Additionally, HCT116 cells pre-incubated with [^13^C_6_]-glucose and then treated with [^18^O_2_]-H_2_O_2_ showed a transient increase in the amount of [^18^O_1_]-acetate peaking around five minutes post induction of ROS with subsequent decay kinetics corresponding to the clearance of the ROS (Fig. 2j)^27^. Importantly, it was observed that up to 30% of the total [^13^C_2_]-acetate pool (Figs. 2j and k) could be derived from a transient increase in ROS from exogenous [^18^O_2_]-H_2_O_2_. This analysis further revealed that acetate is substantially synthesized from pyruvate in cells and the activity of this reaction is mediated by ROS.

While both endogenous and exogenous ROS appear to catalyze in quantitative amounts the conversion to acetate from pyruvate in mammalian cells, the presence of acetate unaccounted by isotopic labeling of oxygen suggests it is not the only mechanism for de novo acetate production. We next tested whether acetate can be released from pyruvate dehydrogenase (PDH) (Fig. 3a). Mammalian PDH mediates oxidative decarboxylation of pyruvate to acetyl-CoA. PDH contains three subunits (E1, E2, and E3). E1 catalyzes pyruvate decarboxylation using the cofactor thiamine pyrophosphate (TPP) and subsequent reductive acetylation of the lipoyl groups. E2 transfers the acetyl group to the thiol group in CoA and generates acetyl-CoA, and E3 regenerates the cofactor lipoate by coupling the oxidation to NAD^+^ reduction. Therefore, the cofactors CoA and NAD^+^ are required for PDH to achieve full activity. However, when CoA levels are lower than pyruvate levels, it may induce limitations at the E2 step. Possibly, other thiol-containing molecules, such as GSH could serve as the acetyl group acceptor, considering that the cellular CoA is around 10 µM while GSH is 1 to 7 mM (Fig. 3b), and similar concentration values in different types of cells or tissues were also reported by others^28–30^. Low levels of CoA and high levels of GSH were also observed in autochthonous mouse tumors providing a physiological context for this occurrence (Fig. 3c). Meanwhile, pyruvate is at much higher concentrations (Figs. 1d and 3b). To study how these imbalanced substrate and cofactor concentrations affect PDH function, we incubated PDH isolated from porcine heart with [^13^C_3_]-pyruvate in the presence or absence of cofactors (TPP, CoA and NAD^+^) (Fig. 3d). Indeed, with CoA at saturated concentrations (400 µM), more than 99% of the pyruvate (200 µM) was consumed in less than 2 mins (Fig. 3d). Surprisingly, in the absence of CoA and NAD^+^, pyruvate was still consumed by PDH in the presence of TPP, but instead converted to [^13^C_2_]-acetate and [^13^C_2_]-acetaldehyde (Figs. 3d, 3e and Supplementary Fig. 3a), with lower overall PDH activity (Fig. 3f). Furthermore, in the absence of CoA, GSH was found to be an acetyl group acceptor and increased overall PDH activity (Fig. 3f) that resulted in the formation of [^13^C_2_]-acetyl-GSH (Ac-GSH) (Fig. 3g). Together these data indicate that when provided with TPP, the mammalian PDH E1 subunit preserves its decarboxylase activity and produces acetaldehyde and acetate in the absence of cofactors CoA and NAD^+^, with reduced overall activity (Fig. 3h). Interestingly, another keto acid dehydrogenase, alpha-ketoglutarate dehydrogenase (aKGDH), that normally catalyzes the conversion from alpha-ketoglutarate to succinyl-CoA via a similar mechanism as PDH, also functions as a pyruvate decarboxylase in the absence of CoA and NAD^+^, catalyzing the conversion of pyruvate to acetaldehyde and to acetate to a smaller extent (Supplementary Fig. 3b).

**Figure 3.**
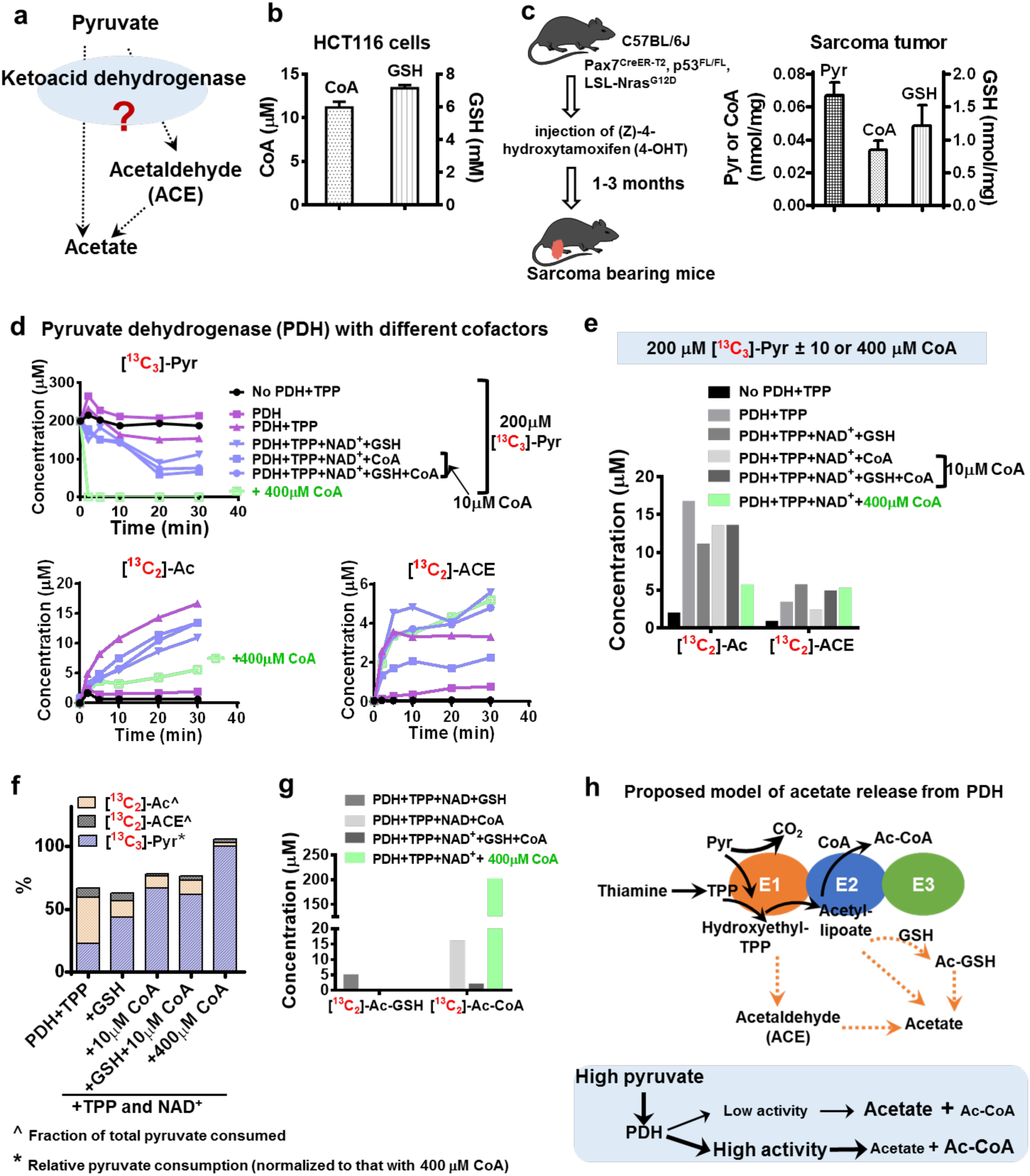
Conversion from pyruvate to acetate is catalyzed by mammalian PDH in a thiamine dependent manner. **a,** Release of acetate and acetaldehyde from pyruvate is possibly catalyzed by keto acid dehydrogenase, especially pyruvate dehydrogenase (PDH). **b**, Intracellular CoA and GSH concentrations in HCT116 cells. **c**, CoA and GSH concentrations in mouse sarcoma tumor. **d**, Conversion of pyruvate (200 µM) to acetate and acetaldehyde by PDH over 30 min in the absence or presence of cofactors. **e**, Concentrations of pyruvate-derived acetate, acetaldehyde after 30 min incubation with PDH. **f**, Pyruvate consumption rate (blue), relative to that in the presence of TPP, NAD^+^ and CoA, representing relative activity of PDH; Acetate (yellow) and acetaldehyde (grey) production, relative to total pyruvate consumption. **g**, Ac-CoA and Ac-GSH production. **h**, Proposed model of acetate release from pyruvate catalyzed by PDH. Error bars obtained from SD of n=3 (HCT116 cells) and n=5 (mouse) independent measurements.

These results strongly suggest that thiamine dependent mammalian keto acid dehydrogenases function as pyruvate decarboxylases when the activities of the E2 and E3 subunits are limited. We then tested whether this also occurred in cultured cells and mice. Indeed, [^13^C_2_]-acetate, [^13^C_2_]-acetaldehyde and [^13^C_2_]-Ac-GSH were quickly formed in HCT116 cells within a few minutes after [^13^C_6_]-glucose medium was added (Fig. 4a) and further confirmed with tandem mass spectrometry (Supplementary Fig. 4a). The intracellular concentrations of [^13^C_2_]-acetate and [^13^C_2_]-acetaldehyde were also found to be comparable to [^13^C_3_]-pyruvate (Fig. 4c). Production of acetate and Ac-GSH was decreased in cells engineered with CRISPR/CAS9 to lack PDH (Fig. 4d and supplementary Figs. 4b and 4c) or when cultured in thiamine depleted media (Fig. 4e). CPI-613, a lipoate analogue and a PDH inhibitor, increased the formation of Ac-GSH (Supplementary Fig. 4d). These results together link PDH activity to acetaldehyde, Ac-GSH, and acetate production, consistent with our findings using purified PDH enzyme (Fig. 3). Interestingly thiamine had a larger effect on acetate generation than complete PDH knockout. This is consistent with pyruvate also being a substrate of other thiamine dependent keto acid dehydrogenases, including alpha ketoglutarate dehydrogenase (Supplementary Fig. 3b) and thiamine depletion decreasing ROS activity as evidenced by decreased ROS-mediated methionine sulfoxide formation (Supplementary Figs. 4e to g).

**Figure 4.**
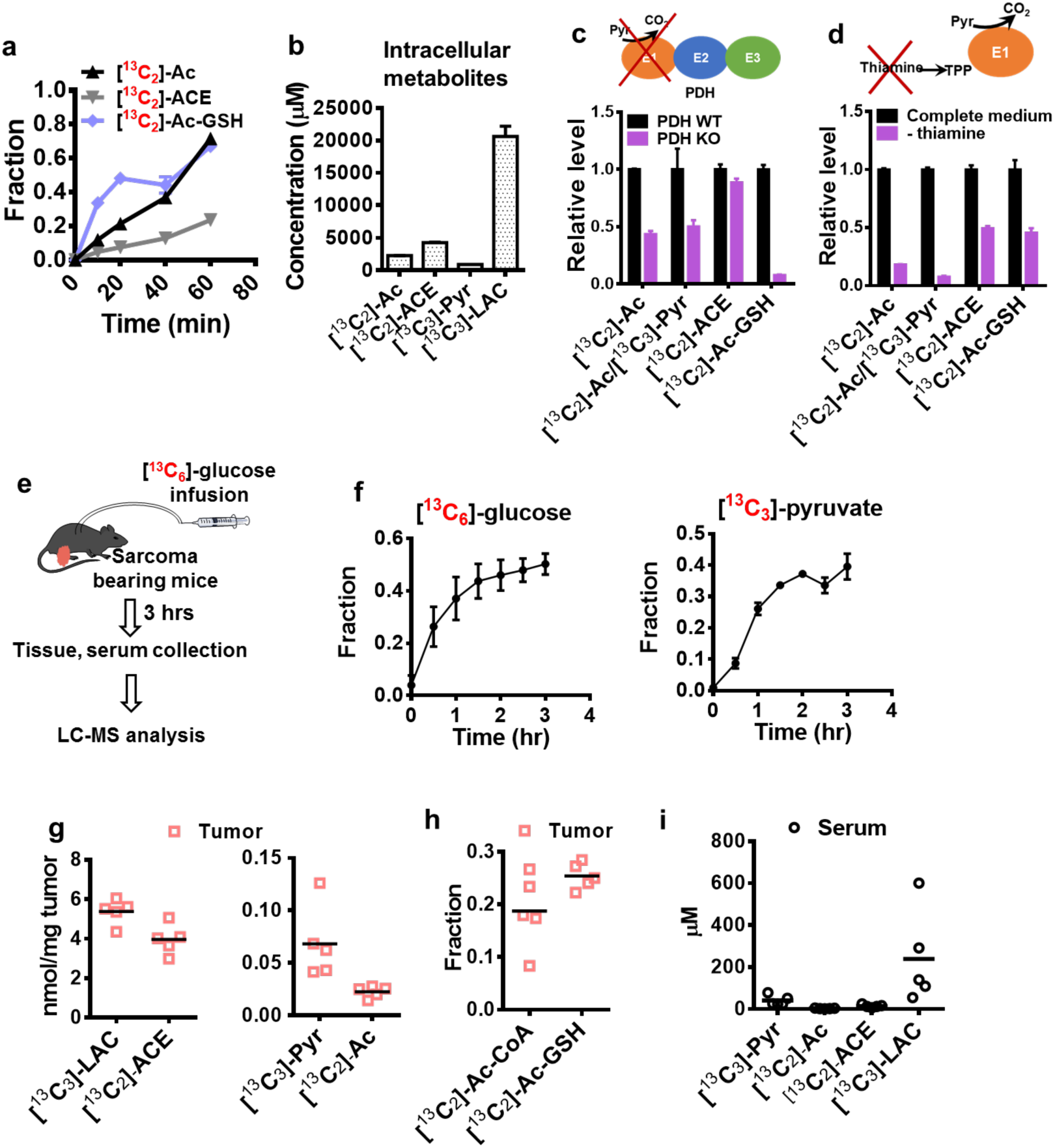
Acetate production from pyruvate occurs in both cultured cells and *in vivo*. **a,** ^13^C enrichment of acetate (Ac), acetaldehyde (ACE) and Acetyl-GSH (Ac-GSH) in HCT116 cultured in the presence of [^13^C_6_]-glucose. **b**, Intracellular concentrations of ^13^C labelled Ac, ACE, Pyr, and LAC in HCT116 cells after 5 hrs incubation in[^13^C_6_]-glucose medium. **c**, Relative level of ^13^C enriched Ac, ACE and Ac-GSH in mouse sarcoma cells with wild type PDH (PDH WT) or PDH knockout (PDH KO) using CRISPR. **d**, Relative level of ^13^C enriched Ac, ACE and Ac-GSH in HCT116 cells cultured in medium with or without thiamine. **e**, Scheme of ^13^C glucose infusion in sarcoma bearing mice. **f**, ^13^C enrichment fraction of glucose and pyruvate in the serum of sarcoma free mouse after the [^13^C_6_]-glucose infusion. **g**, Concentrations of [^13^C_3_]-Pyr, [^13^C_3_]-LAC, [^13^C_2_]-ACE and [^13^C_2_]-Ac in sarcoma tumor. **h**, ^13^C enrichment of Ac-GSH and [^13^C_2_]-Ac-GSH in sarcoma. **i**, Concentrations of [^13^C_3_]-Pyr, [^13^C_3_]-LAC, [^13^C_2_]-ACE and [^13^C_2_]-Ac in sarcoma tumor. **g to i** were from sarcoma mouse samples collected 3 hrs after [^13^C_6_]-glucose infusion. Error bars obtained from SD of n=3 (HCT116 cells) and n=5 (mouse) independent measurements.

To determine whether these reaction mechanisms could occur in a physiological context, we first established a euglycemic transfusion system where mice bearing KRAS^+/G12D^; TP53^-/-^ autochthonous sarcoma tumors (Fig. 3c) were continuously infused [^13^C_6_]-glucose^31,32^ (Fig. 4e). We first conducted this experiment using tumor-free mice, and observed that over 40% of the glucose in the serum at steady state could be labeled [^13^C_6_]-glucose resulting in approximately 35% [^13^C_3_]-pyruvate labeling (Fig. 4f). Next in tumor-bearing mice, both serum and tumor tissue were collected for metabolite analysis after 3 hrs of glucose infusion. ^13^C labeled pyruvate, lactate, acetate and acetaldehyde were detected in tumor tissue at different concentrations (Fig. 4g). The enrichment of ^13^C labelled [^13^C_2_]-Ac-CoA and [^13^C_2_]-Ac-GSH was comparable (Fig. 4h). The concentrations of [^13^C_3_]-pyruvate, [^13^C_3_]-lactate and [^13^C_2_]-acetate were much higher in the tumor than in the serum (Fig. 4i), indicating that these metabolites were produced locally confirming our findings in vitro and in cultured cells (Fig. 3 and Figs. 4a–d).

To expand the scope of the study and thus identify contexts in which these pathway is biologically necessary, we sought to explore contexts in which acetate derived from some cells could interact with the metabolism of other cells. ATP citrate lyase (ACLY) is a cytosolic enzyme that converts nucleocytosolic citrate to Ac-CoA and acetate requirements in cells increase when its activity is impaired^33^. We thus considered mouse embryonic fibroblasts (MEFs), known to produce low amounts of acetate (Fig 1c) that were engineered to lack ACLY (ACLY KO) (Fig. 5a and 5b) that were previously described and further confirmed to exhibit an acquired dependence on acetate^33^. Co-culturing these MEFs with acetate-producing HCT116 cells separated by a 0.4 micron membrane allowing for free diffusion (Fig. 5c) resulted in dramatically increased medium acetate concentrations in the MEF compartment (Fig. 5d), and rescued deficiencies in the growth rate in the knockout cells (Fig. 5e) with negligible effects on the control (ACLY WT) MEFs (Fig. 5e). Increased de novo lipogenesis indicated by increased incorporation of [^13^C_6_]-glucose into palmitate in ACLY KO cells co-cultured with HCT116 cells confirmed increased cytosolic Ac-CoA production (Fig. 5f). These findings demonstrate that endogenous de novo acetate produced through overflow metabolism from one cell type can be utilized to support proliferation of surrounding cells with acetyl-CoA insufficiencies.

**Figure 5.**
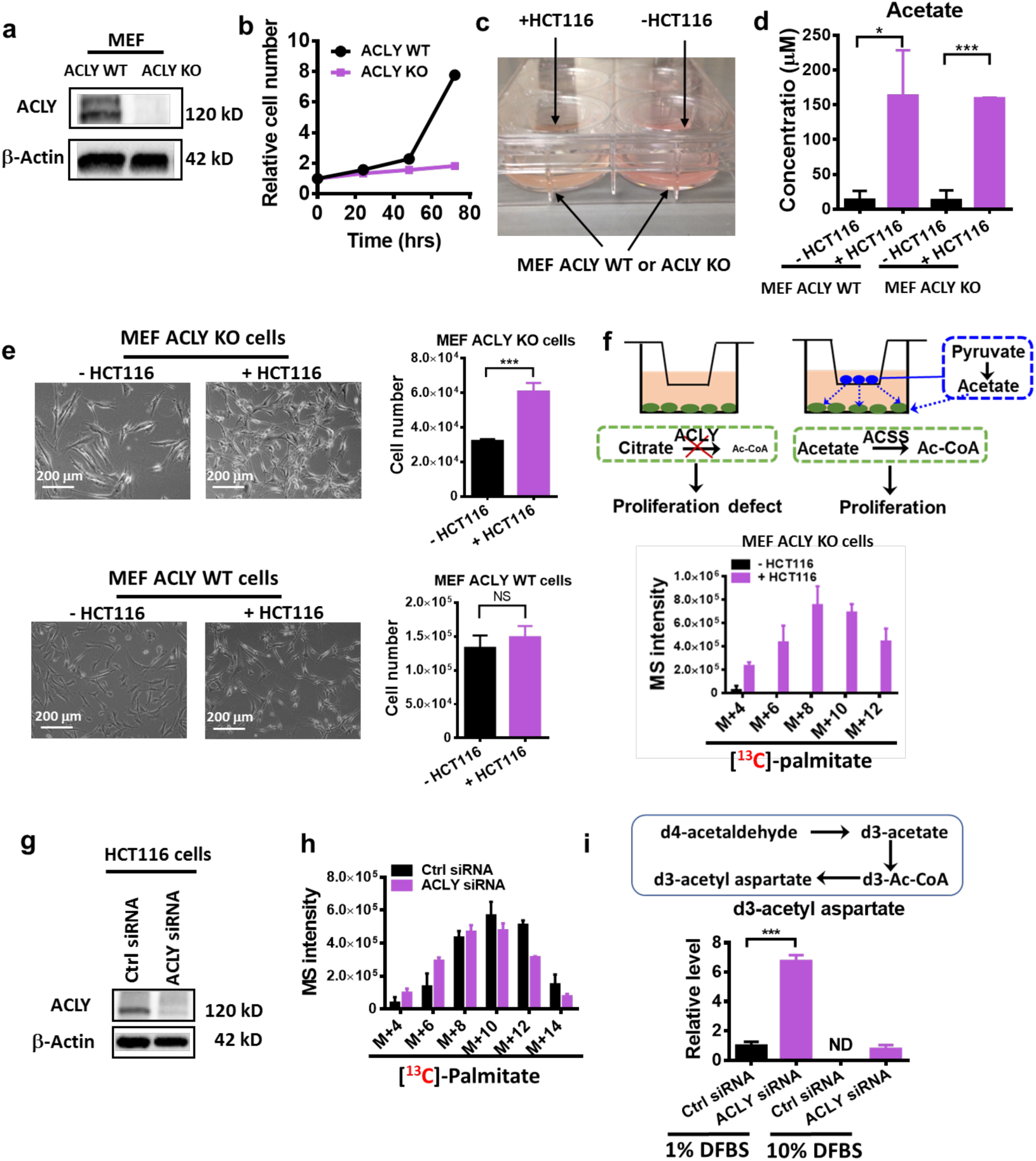
Acetate released rescues cells with ACLY deficiency. **a,** Western blot of ATP citrate lyase (ACLY) in MEF (ACLY WT or KO) cells. **b**, Growth curve of MEF (ACLY WT or KO) cells cultured in RPMI 1640 supplemented with 10% dialyzed FBS. **c**, Photo of co-culture system. **d**, Medium acetate concentration in co-culture system in the absence or presence of HCT116 cells. **e**, Representative photos and cell number of MEF ACLY WT or KO cells co-cultured with or without HCT116 for 3 days. **f**, Model of proliferation rescue of MEF ACLY KO cells by acetate released from HCT116 cells in a co-culture system (top), and incorporation of ^13^C into palmitate in MEF ACLY KO cells co-cultured with or without HCT116 cells in RPMI medium containing [^13^C_6_]-glucose and 10% dialyzed FBS (bottom). **g**, Western blot of ATP citrate lyase (ACLY) in HCT116 cells treated with or without non-targeted control (Ctrl) siRNA or ACLY siRNA. **h**, Incorporation of ^13^C into palmitate in HCT116 cells cultured in RPMI medium containing [^13^C_6_]-glucose and 10% dialyzed FBS for 1hr. **i**, Carbon incorporation into acetyl group from acetaldehyde by HCT116 cells cultured in low (1%) or regular (10%) D-FBS for 1hr. Error bars obtained from SD of n=3 independent measurements.

To further explore this pathway, we examined the effects of ACLY expression in high-acetate producing HCT116. ACLY knockdown had small effects on HCT116 cell de novo lipogenesis (Fig. 5g and 5h) and isotope tracing with deuterated acetaldehyde confirmed that ACLY knockdown cells exhibited increased acetate utilization which could be further increased by limiting nutrient availability during serum starvation (Fig. 5i), which is consistent with previous findings^10,34^. Thus these experiments confirm that de novo pyruvate-derived acetate production can have critical functional roles in supporting acetyl-CoA metabolism.

## Discussion

The dependence on acetate to supply both nucleocytosolic and mitochondrial acetyl-CoA pools has been identified in multiple cell types, including cancer cells, T cells, and neurons^8–8,13,14^. Our study provides evidence for a glucose-derived overflow pathway for acetate metabolism that occurs in mammalian cells. It further identifies two mechanisms with differential regulatory capacities involving non-enzymatic and enzymatic pyruvate-derived acetate chemistry resulting in endogenous acetate production as a new component of mammalian central carbon metabolism.

Acetate generation was shown to in part to be regulated by ROS, a finding which potentially links this pathway to numerous physiological and pathophysiological processes ^35–37^. ROS levels are dynamic and at steady state, intracellular concentrations of H_2_O_2_ vary from nM to µM ^38,39^, while extracellular concentrations of H_2_O_2_ may be 10 to 100 times higher under certain circumstances, such as inflammation^40^. Consistent with this pathway, cancer cells have been shown to produce H_2_O_2_ at a rate of around 1 nmol/10^4^Cells/hour ^41,42^, while the combined pyruvate generation and secretion rate (Fig. 1b) was found to be around 2 nmol/10^4^Cells/hour. Thus, oxidative stress and the activity of the electron transport chain leading to ROS would allow for coupling of acetate production to all metabolic functions within central carbon metabolism ^43,44^. This pathway therefore may allow cells to regulate acetyl-CoA metabolism to some extent during oxidative stress as occurs, for example, during obesity and related sequelae and in nutrient-restricted tumor microenvironments. In these situations, increased ROS levels that can occur through defects in mitochondrial function or exposure to DNA damage results in the production of pyruvate-derived acetate that can be used for lipogenesis, protein acetylation, and energy metabolism.

The other reaction mechanism found to generate acetate occurred through altered enzyme activity of keto-acid dehydrogenases which transformed their activity to keto-acid decarboxylases. The extent of this mechanism appeared to be regulated by the supply of cofactors, such as CoA, and is coupled to the glycolytic rate, mitochondrial TCA cycle activity, and availability of glutathione which is also coupled to oxidative status. Thus, numerous possibilities may exist by which acetate and resulting acetyl-CoA generation is coupled to other critical metabolic functions.

## Acknowledgements

We acknowledge support from National Institutes of Health awards R01CA193256, R00 CA16899 to JWL, T32 CA093240 to DEC, R35 CA197616 to DGK, R01CA174761 to KEW, and F99CA222741 to SZ, the American Cancer Society (TBE434120 to JWL and TBE130927 to MAR), and the King Abdullah International Medical Research Center under the Ministry of National Guard Health Affairs (AAC). We thank Dr. Anthony Ribeiro (NMR facility at Duke University) for help with spectroscopy measurements. We thank Dr. Clementina Mesaros (University of Pennsylvania) for helpful discussions on the ^18^O_2_ tracing assay. We also thank members of the Locasale Lab for helpful discussions.

## Author Contributions

JWL and XL conceived the study, performed all data analysis and interpretation, and wrote the paper. Unless noted, XL performed all experiments with pilot studies performed by AAC and MOW. XL and JWL wrote the paper. DEC and DGK designed and performed the animal experiments, shared sarcoma cell lines, interpreted data and edited the paper. SZ and KEW generated the MEF (ACLY WT and KO) cell lines, provided advice on assay development, and edited the paper. MAR helped interpreting data and edited the paper. JL helped to establish the ^18^O_2_ tracing assay.

---

**Supplementary Figure 1.**
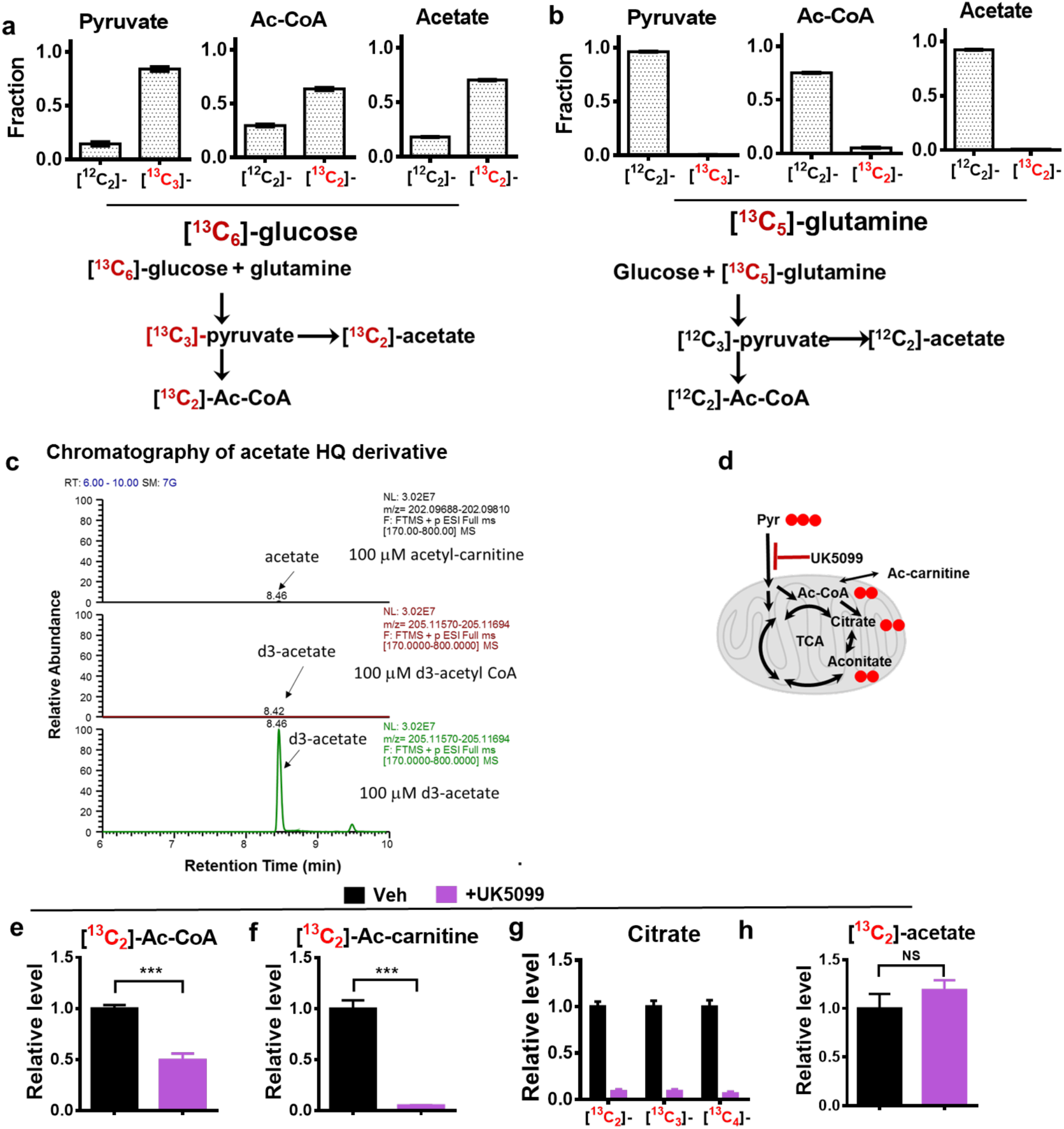
Evaluation of glutamine and pyruvate carrier inhibition contribution to acetate and TCA metabolites. a, ^13^C enrichment pattern of acetate released from HCT116 cells treated with [^13^C_5_]-glutamine or [^13^C_6_]-glucose. **b**, Extracted ion chromatography of acetate or d3-acetate HQ derivative from samples prepared by incubating acetyl-carnitine, d3-acetyl-CoA or d3-acetate (100 µM) with HQ derivatization reagent at 37 °C for 70 min. c, ^13^C enrichment patterns of metabolites derived from [^13^C_6_]-glucose. Red solid circle denotes ^13^C. Ac: Acetate; Ac-carnitine: Acetyl-carnitine; Pyr: Pyruvate. The relative level of ^13^C enriched Ac-CoA (a), Ac-carnitine (b), citrate (c), and acetate (d) in HCT116 cells cultured in RPMI 1640 medium containing [^13^C_6_]-glucose with or without UK5099 treatment for 40 min. Error bars obtained from SD of n=3 independent measurements.

**Supplementary Figure 2.**
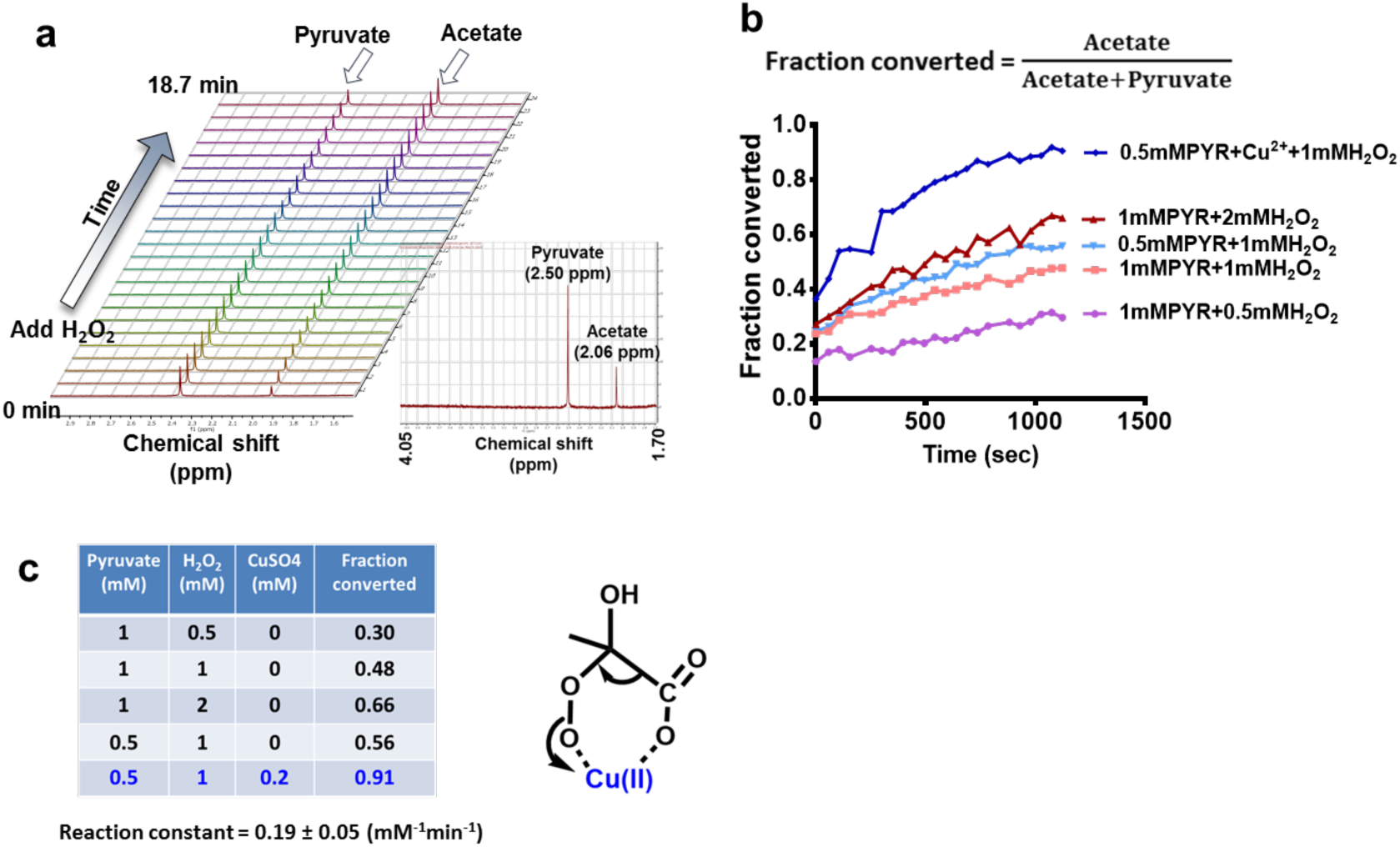
Exogenous H_2_O_2_ catalyzes pyruvate decarboxylation to acetate. **a**, Representative NMR spectrum for pyruvate and acetate peaks from time 0 to 18.7 min. Time 0 min is when data acquisition began, and there is a delay from actual reaction time due to NMR tube temperature equilibration. **b**, Conversion rate of pyruvate (Pyr) to acetate under different conditions from 0 to 18.7 min since the first data acquisition. Conversion rate was calculated by dividing acetate peak area by the sum of acetate and pyruvate peak area. **c**, Summary of reaction conditions, conversion rate (at 18.7 min), and reaction constant (mean ± SD).

**Supplementary Figure 3.**
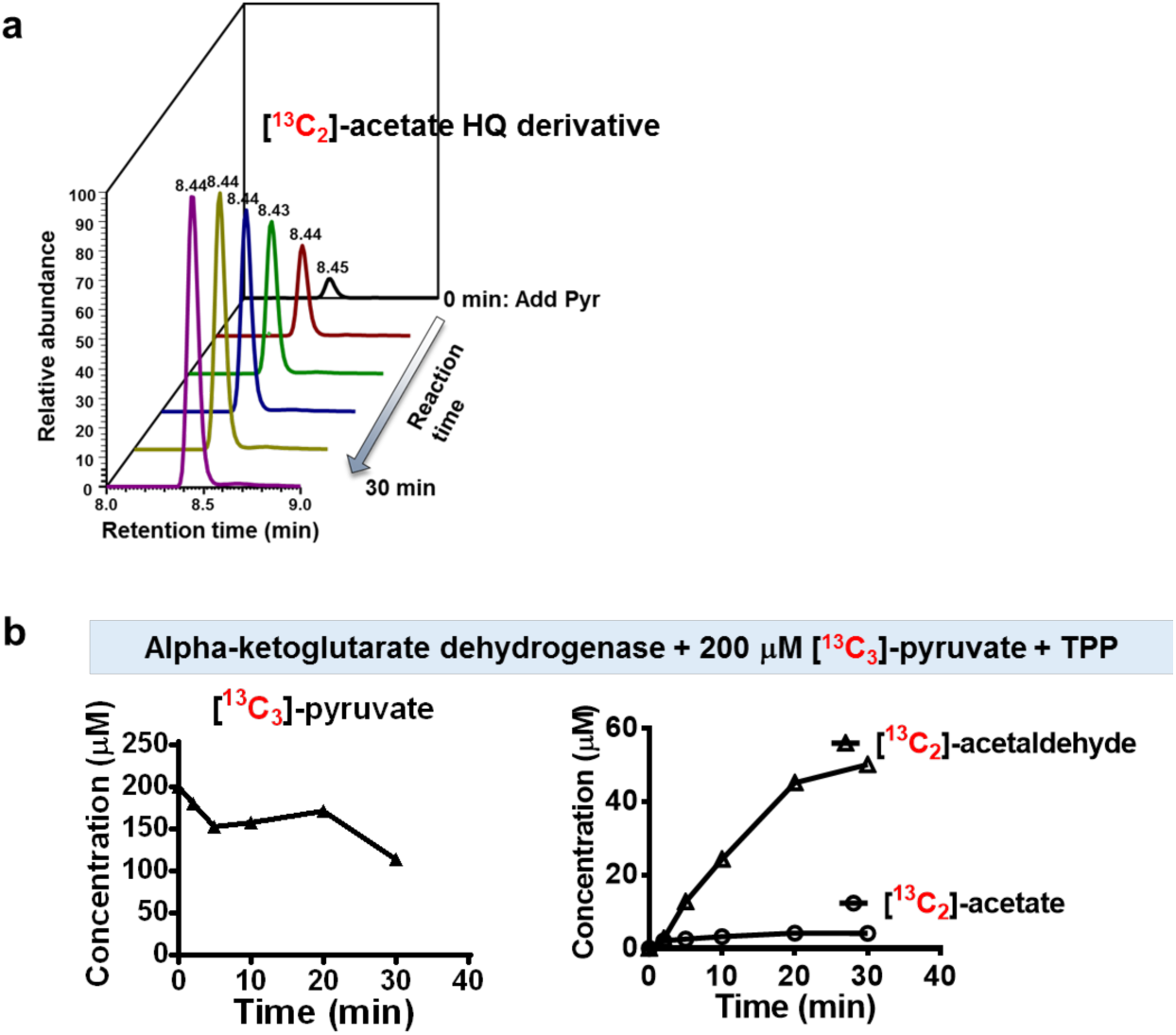
Keto-acid dehydrogenases catalyze acetate and acetaldehyde production from pyruvate in the absence of cofactors CoA and NAD^+^. **a**, Release of acetate from pyruvate in the presence of pyruvate dehydrogenase (PDH) supplemented with thiamine pyrophosphate (TPP). **b**, Consumption of pyruvate and production of acetaldehyde and acetate by alpha-ketoglutarate dehydrogenase supplemented with TPP.

**Supplementary Figure 4.**
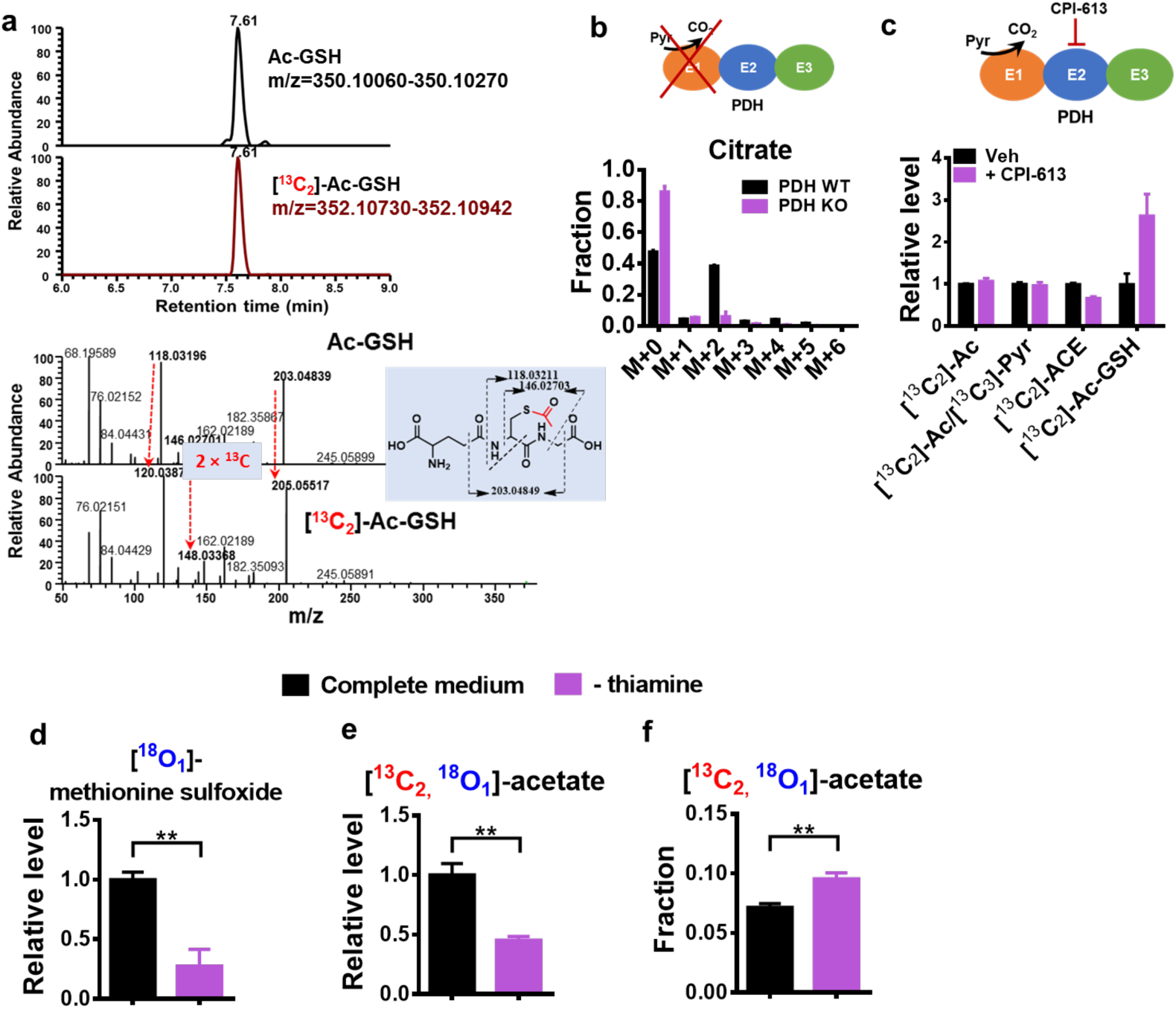
Metabolites from HCT116 cells subjected to alterations of mitochondrial metabolism. **a**, Extracted ion chromatography and MS2 spectrum (positive ion mode) of [^13^C_2_]-Ac-GSH in HCT116 cells cultured in [^13^C_6_]-glucose medium for 40 min. **b**, ^13^C enrichment of citrate in mouse sarcoma (PDH WT and KO) cells cultured in ^13^C glucose for 6 hrs. **c**, Relative level of ^13^C enriched Ac, ACE and Ac-GSH in HCT116 cells in the absence of presence of CPI-613, a lipoate analog. **d to f**, The effect of thiamine depletion on the formation of [^13^C_2_, ^18^O_1_]-Ac, [^18^O_1_]-methionine sulfoxide in HCT116 cells cultured in the presence of ^18^O_2_ for 48 hrs. For thiamine depletion, HCT116 cells were cultured in thiamine free medium for 4 days before ^18^O_2_ tracing. Error bars obtained from SD of n=3 independent measurements.

## Methods

### Reagents

RPMI 1640 medium was purchased from Cellgro. Thiamine free RPMI 1640 was prepared in house. Fetal Bovine Serum (FBS) were purchased from Hyclone Laboratories. Dialyzed FBS (DFBS) was obtained from ThermoFisher Scientific. Optima ammonium acetate, ammonium hydroxide, Optima LC-MS grade, acetonitrile, methanol and water were purchased from Fisher Scientific. [^13^C_6_]-D-glucose, d3-sodium pyruvate and [^13^C_3_]-sodium pyruvate were obtained from Cambridge Isotope Laboratories. Sodium pyruvate, sodium lactate, potassium acetate, sodium chloride, UK5099, copper sulfate, 2-hydrazinoquinoline, triphenylphosphine, 2,2′-dipyridyl disulfide, D_2_O, [^18^O_2_]-H_2_O_2_ (2–3% solution), ^18^O_2_, d4-acetaldehyde, d3-sodium acetate, ammonium tetrathiomolybdate (TTM), CPI-613, RPMI 1640 (without sodium bicarbonate), HEPES buffer, pyruvate dehydrogenase and ketoglutarate dehydrogeanse (from porcine heart) were purchased from Sigma. d3-acetyl-carnitine hydrochloride was purchased from Santa Cruz Biotechnology. Balloons were purchased from local stores. Mini vacuum desiccator (Bel-Art) was purchased from VWR.

### Cell culture

MEF (ACLY WT (*Aclyf/f*) and KO) cells were prepared as described previously^33^. HCT116, SKOV3, HepG2, and BT474 cell lines were obtained from ATCC. The HeyA8 cell line was a generous gift from Dr. Ernst Lengyel’s lab, University of Chicago, IL. All human cell lines used in the study were authenticated at Duke University DNA Analysis Facility using GenePrint 10 kit from Promega and tested to be mycoplasma-free. All cell culture related medium and reagents were sterilized using 0.22 µm sterile filter. Mouse sarcoma cells were derived from sarcoma bearing mice, and the procedure is described below.

All cells were first cultured in 10 cm dish with full growth medium (RPMI 1640 supplemented with 10% FBS). The cell incubator was set at 37 °C supplemented with 5% CO_2_. For cell proliferation assay, cells were cultured in 12 well plate, and in the end of treatment, cells were first harvested by trypsinization, and cell number and volume were measured using Moxi™ Z Mini Automated Cell Counter (ORFLO Technologies). For metabolite analysis, cells were seeded into 6 well plate at the density of 500 000 cells per well. After overnight incubation in full growth medium, for UK5099 treatment, the old medium was removed and cells were briefly washed with glucose free RPMI 1640 before 1.5 ml of RPMI 1640 (supplemented with 10% dialyzed FBS) with 0.025% DMSO or 5 µM UK5099 (final DMSO content 0.025%) was added to each well. After 5 hours incubation, medium was replaced with 1 ml of RPMI 1640 containing 11.1 mM [^13^C_6_]-glucose, 10% dialyzed FBS, and 0.025% DMSO or 5 µM UK5099 (final DMSO content 0.025%). For acetate measurement, 5 µl medium from each well was collected at different time points and immediately placed on dry ice before storage in -80 °C freezer. Intracellular metabolites were extracted at 10, 20 and 40 min, and the detailed extraction procedure will be described below. For [^18^O_2_]-H_2_O_2_ treatment, after incubation in [^13^C_6_]-glucose medium, [^18^O_2_]-H_2_O_2_ was directly added to each selected well (final concentration, 10, 30 or 300 µM). 5 µl medium from each well was collected at 5, 10, 20 and 40 min. Intracellular metabolites were extracted at 10 min after addition of [^18^O_2_]-H_2_O_2_ (300 µM). For CPI-613 treatment, HCT116 cells were incubated in RPMI 1640 (supplemented with 10% dialyzed FBS) in the absence or presence of 150 µM CPI-613 for 4 hours and media were replaced with 1 ml of RPMI 1640 containing 11.1 mM [^13^C_6_]-glucose, 10% dialyzed FBS, and 0.15% DMSO or 150 µM CPI-613 (final DMSO content 0.15%). Media were collected after 40 min, and intracellular metabolites were analyzed after 3-hr incubation in ^13^C-glucose media. For thiamine deprivation assay, HCT116 cells were incubated in thiamine free RPMI 1640 (supplemented with 10% dialyzed FBS) for 4 days and media were replaced with 1 ml of thiamine free RPMI 1640 containing 11.1 mM [^13^C_6_]-glucose and 10% dialyzed FBS. Media and intracellular metabolites were collected after 40-min incubation in ^13^C-glucose media.

### Animal

All animal procedures for this study were approved by the Institutional Animal Care and Use Committee (IACUC) at Duke University. 8 to 10-week old C57BL/6 J mice (Jackson Laboratory) were used to develop the ^13^C-glucose infusion workflow. Mouse models of soft tissue sarcoma were generated in a mixed 129/SVJae and C57BL/6 background (Jackson Laboratory) using a combination of alleles: Pax7^CreER-T2^, p53^FL/FL^, LSL-Nras^G12D^. Primary mouse soft tissue sarcomas were generated in the mouse hind limb by intramuscular (IM) injection of (Z)-4-hydroxytamoxifen (4-OHT). A euglycemic glucose infusion was performed according to a procedure previously described ^32^. Briefly, a jugular vein catheter was surgically implanted and exteriorized via a vascular access port, which allows infusion of [^13^C_6_]-glucose via the venous catheter. Blood from a tail nick in freely-moving and unanesthetized animals was serially withdrawn. Four to five days after surgery mice were fasted for 6 hours and continuously infused with [^13^C_6_]-glucose at a rate of 20 mg/kg/min for a total of 3 hours. Blood was collected every 30 min for the duration of the infusion. Immediately following infusion, animals were sacrificed and liver tissue was excised and rapidly frozen in liquid N_2_ to quench metabolism. Tissue was stored at -80°C until used for metabolite extractions. No data were excluded from the analysis. For these genetically engineered mice, no comparison was made between multiple groups, so randomization is not applicable.

### siRNA transfections

siRNA transfection was performed following manufacturer’s instructions with Lipofectamine RNAiMAX (Invitrogen), using siRNA pools targeting human ACL (Dharmacon #L- 004915–00), or a non-targeting control (Dharmacon #D-001810–10-20) at a concentration of 20 nM in growth medium (RPMI 1640 with 10% FBS). After 48 hrs, cells were split into 6 well plate. All analysis (protein, metabolite and cell proliferation) was performed after 72 hrs of siRNA transfections.

### Western blotting

Cells were collected into ice cold RIPA buffer containing cOmplete™ Protease Inhibitor Cocktail (Roche). Protein lysate was first loaded to a 12% Mini-PROTEAN TGX Precast Protein Gels (Bio-Rad) for separation, and then transferred to nitrocellulose membrane using Trans-Blot SD Semi-Dry Transfer Cell (Bio-Rad). The blot was visualized using ChemiDoc (Bio-Rad). ATP citrate lyase antibody (rabbit, 15421–1-AP) was purchased from Proteintech. β-actin antibody (mouse, 8H10D10) was purchased from Cell signaling. Anti-mouse IgG (goat, 610–1102) and anti-rabbit IgG (donkey, 611–7302) antibody were obtained from Rockland Immunochemicals Inc.

### HPLC method

Ultimate 3000 UHPLC (Dionex) was used for metabolite separation and detection. For polar metabolite analysis, a hydrophilic interaction chromatography method (HILIC) with an Xbridge amide column (100 x 2.1 mm i.d., 3.5 µm; Waters) was used for compound separation at room temperature. The mobile phase and gradient information were described previously ^45^. 2-hydrazinoquinoline derivatives were measured using reversed phase LC method, which employed an Acclaim RSLC 120 C8 reversed phase column (150 x 2.1 mm i.d., 2.2 µm; Dionex) with mobile phase A: water with 0.5% formic acid, and mobile phase B: acetonitrile. Linear gradient is: 0 min, 2% B; 3 min, 2% B; 8 min, 85% B;9.5 min, 98%B; 10.8 min, 98%B, and 11 min, 2% B. Flow rate: 0.2 ml/min. Column temperature: 25 °C.

### Mass Spectrometry

The Q Exactive Plus mass spectrometer (HRMS) is equipped with a HESI probe, and the relevant parameters are as listed: heater temperature, 120 °C; sheath gas, 30; auxiliary gas, 10; sweep gas, 3; spray voltage, 3.6 kV for positive mode and 2.5 kV for negative mode. Capillary temperature was set at 320 °C, and S-lens was 55. A full scan range was set at 70 to 900 (*m/z*) with positive/negative switching when coupled with the HILIC method, or 170 to 800 (*m/z*) at positive mode when coupled with reversed phase LC method. The resolution was set at 140 000 (at *m/z* 200). The maximum injection time (max IT) was 200 ms at resolution of 70 000 and 450 ms at resolution of 140 000. Automated gain control (AGC) was targeted at 3 × 10^6^ ions. For targeted MS2 analysis, the isolation width of the precursor ion was set at 1.0 (*m/z*), high energy collision dissociation (HCD) was 35%, and max IT is 100 ms. The resolution and AGC were 35 000 and 200 000, respectively.

### 2-hydrazinoquinoline derivatization

To measure acetate, pyruvate, lactate and acetaldehyde, we developed a method based on previous publication ^47^. 20 mM of 2-hydrazinoquinoline (HQ), triphenylphosphine, and 2,2′-dipyridyl disulfide (DPDS) were freshly prepared in acetonitrile solution, except that for DPDS, methanol (final concentration, 2.5%) was added to facilitate dissolution. HQ derivatization reagent cocktail in acetonitrile (0.5 mM of HQ, Triphenylphosphine, and DPDS) was then prepared from 20 mM stock solution. 2 µl of cell culture medium, mouse serum or intracellular metabolite extract in 80% methanol was added to 1.5 ml Eppendorf tube containing 200 µl of 0.5 mM HQ derivatization reagent cocktail in acetonitrile. d3-pyruvate, d3-acetate, and d3-lacate were used as internal standards to facilitate absolute quantification. After rigorous vortexing, the samples were incubated in water bath at 37 °C for 70 min. After derivatization, the samples were then centrifuged at 4 °C for 10 min at the speed of 20 000 g. The supernatant was then added formic acid (final concentration, 1%) and transferred to a new LC vial before LC-HRMS analysis. To check potential artifact, 2 µl 100 µM d3-acetyl-CoA or acetyl-carnitine was incubated with the HQ derivatization reagents (Supplementary Fig. 1c). In addition, LC-MS grade water, 80% MeOH, and cell free medium were always included to check the baseline level of acetate, pyruvate, lactate and acetaldehyde. For acetate and acetaldehyde measurement, evaporation process was avoided due to the low boiling point of acetaldehyde.

### Metabolite extraction from cultured cells

For intracellular metabolite measurement, at 10, 20 and 40 min, the medium was removed, cells were briefly washed twice with 1 ml ice cold saline solution (0.9% NaCl in water) before placed on dry ice, followed by the addition of 1 ml 80% methanol/water (pre-cooled in -80 °C freezer) to each well. After incubation in -80 °C freezer for 15 min, cells were scraped into 80% methanol on dry ice and then transferred to Eppendorf tube. Samples were centrifuged at 20 000 g for 10 min at 4 °C, and the supernatant was split into two Eppendorf tubes before drying in speed vacuum concentrator (Labcono). For absolute quantitation of GSH and CoA, ^13^C labelled E.coli metabolite extract^46^ was added to the cell metabolite extract before drying. The dry pellets were reconstituted into 30 µl sample solvent (water:methanol:acetonitrile, 2:1:1, v/v) and 3 µl was injected into the LC-HRMS. The sample sequence of data acquisition was randomized.

### Metabolite extraction from tissue

Briefly, the tumor sample was first homogenized in liquid nitrogen and then 15 to 25 mg was weighed in a 2 ml vial pre-filled with Zirconia/Silica beads (1.0 mm dia). Extraction solvent 80% methanol/water (pre-cooled in -80 °C freezer) was added to each sample at the ratio of 100 µl/15 mg on dry ice, and a Beadbeater (Biospec) was used to facilitate the metabolite extraction at shaking speed of 3400 rpm for 1 min at room temperature. The metabolite extract was then immediately centrifuged with the speed of 20 000 g at 4 °C for 10 min. The supernatant of tumor extract collected from sarcoma bearing mice which received ^13^C-glucose infusion were used for polar metabolite measurement. For absolute quantitation of GSH and CoA, ^13^C labelled E.coli metabolite extract^46^ was added to the supernatant of tumor extract collected from sarcoma bearing mice without ^13^C-glucose infusion and 3 µl was directly injected to LC-HRMS.

### ^18^O_2_ tracing assay

Cells were seeded into 35 mm dish at the density of 80 000 (SKOV3) or 120 000 (HCT116) in growth medium (RPMI 1640 supplemented with 10% FBS). Right before ^18^O_2_ tracing assay, the medium was replaced with 2 ml RPMI 1640 (without NaHCO_3_), containing 11.1 mM [^13^C_6_]-glucose, 20 mM HEPES and 10% dialyzed FBS (DFBS). For drug treatment, TTM (final: 25 µM) or UK5099 (final: 5 µM) was added to the medium. Cells in 35 mm dishes were then transferred to a 500 ml container (desiccator) containing 1 ml sterile water to maintain the humidity inside the container. The container was subjected to vacuum for 15 seconds and then immediately filled with 100 ml ^18^O_2_ and 400 ml N_2_. The container was then sealed with a 3-way valve and large paper clips were also employed to ensure air tightness. Balloons were employed for gas transfer and to balance the gas pressure in the container. The sealed container was then placed in a non-CO_2_ incubator set at 37 °C. Here, HEPES-buffered medium was used to replace CO_2_/HCO_3_ buffer system to prevent the medium pH fluctuation during the 15-sec vacuum treatment. In the end of treatment (15, 48, or 60 hrs), the spent media were collected and immediately placed on dry ice. Intracellular metabolites were harvested as described previously.

### Co-culture assay

Transwell^®^ cell culture inserts (Corning, Polycarbonate, pore size: 0.4 µm) were used to set up the co-culture system. Briefly speaking, MEF (ACLY WT and KO) cells were seed into 6 well plate at the density of 25 000/well, and HCT116 cells were seeded into the corresponding insert at the density of 50 000/insert. After overnight incubation in growth medium, the medium was replaced with 2.5 ml RPMI 1640 supplemented with 10% DFBS. The co-culture plates were then placed back to CO_2_ incubator. After 48 hrs, cell number was counted using (Moxi Z Mini Automated Cell Counter, ORFLO Technologies). Medium was collected for acetate measurement, and intracellular metabolites were collected for palmitate measurement.

### Purified pyruvate dehydrogenase assay

Pyruvate dehydrogenase (PDH) from porcine heart (Sigma, P7032-10UN) and alpha-ketoglutarate dehydrogenase (aKGDH) from porcine heart (Sigma, K1502-20UN) were supplied as a 50% glycerol solution containing ~9 mg/mL bovine serum albumin, 30% sucrose, 1.5 mM EDTA, 1.5 mM EGTA, 1.5 mM 2-mercaptoethanol, 0.3 TRITON^®^ X-100, 0.003% sodium azide, and 15 mM potassium phosphate, pH 6.8. PDH or aKGDH solution was diluted to 1 unit/ml with buffer containing 20 mM sodium phosphate and 1 mM MgCl_2_, pH7.2. Mixture of thiamine pyrophosphate (TPP, 50 µM), NAD^+^ (5 mM), glutathione (GSH, 40 mM) and Coenzyme A (CoA, 1 mM) were freshly prepared in the same buffer and diluted by 10 times before use. Reaction buffer in Eppendorf tube (without [^13^C_3_]-pyruvate) was incubated in 37 °C water bath for 1 min, and the reaction was initiated by adding [^13^C_3_]-pyruvate (final concentration: 200 µM). For acetate, acetaldehyde, and pyruvate measurements, at 0, 2, 5, 10, 20 and 30 min, 2 µl reaction mixture was collected and immediately added to 200 µl HQ solution in acetonitrile. HQ derivatization was performed as described previously. At 30 min, the reaction was quenched by adding 4 volumes of ice cold methanol. Acetyl-CoA was also added as an internal standard to quantify [^13^C_2_]-acetyl-CoA generated from PDH reaction. After vigorous vortexing, the mixture was centrifuged at 20 000 rcf for 10 min at 4 °C. The supernatant was directly used for acetyl-CoA and acetyl-GSH analysis by LC-HRMS.

### Generation of mouse sarcoma cell lines and CRISPR-based PDH knockout cell lines

Mouse primary sarcoma cell lines were generated from Pax7^CreER-T2^, p53^FL/FL^, LSL-Nras^G12D^ tumors as described previously^48^. Briefly, tumor tissue was excised and digested with 5 mg/mL Collagenase Type I at 37 °C for 45 mins. Red blood cells were lysed and cell pellets washed with PBS before plating. Cells were passaged 4–5 times to deplete stromal cells. PDH knockout cell lines were generated using lentiviral transduction of primary cells with lentiCRISPR v2 backbone, which contains mammalian Cas9, a sgRNA for PDH (5-catgccatagcggttgttct-3’), and puromycin resistance marker. LentiCRISPRv2 was a gift from Feng Zhang (Addgene plasmid # 52961)^49^. 48 hours after transduction, cells were treated with 5 ug/uL puromycin to select for infected cells. Once selected, cells were diluted and plated to sort single-cell colonies. Single-cell outgrowth was confirmed by sequencing and TIDE analysis^50^. Loss of PDH activity was confirmed by culturing PDH WT and KO cells in RPMI medium containing [^13^C_6_]-glucose for 6 hrs. ^13^C metabolite analysis was performed on LC-HRMS as described above. All LC-HRMS analysis was conducted in a blinded fashion.

### Nuclear Magnetic Resonance kinetics assay

An NMR tube containing 0.6 ml of D_2_O was pre-warmed at 37 °C and to check the background, 6 µl of freshly prepared pyruvate stock solution (50 mM or 100 mM in D_2_O) was added. NMR tube was then equilibrated to 37 °C before data acquisition on 500 MHz Varian Inova spectrometer. After first 8 scans, 6 µl of H_2_O_2_ stock (50, 100, or 200 mM in D_2_O) was added to NMR tube to initiate the reaction. A further equilibration of reaction mixture in NMR tube to 37 °C was required before data acquisition. To check the transition metal effect on this reaction, CuSO4 (final 200 µM) was added to NMR tube before the addition of H_2_O_2_. After temperature equilibration, data was then collected continuously for 24 data points (8 scans per data point, roughly 48 seconds per data point). However, it is worth mentioning that due to instrument limitations, the first data point collected had a delay from the real reaction time, and we performed the reaction at 37 °C to mimic the physiology temperature.

### Data analysis

LC-MS peak extraction and integration were performed using commercial available software Sieve 2.0 (ThermoFisher Scientific). The integrated peak area was used to calculate the fold changes between different treatments and ^13^C or ^18^O enrichment. All data are represented as mean ± SD. All p values are obtained from student’s t-test two-tailed using GraphPad Prism 6 unless otherwise noted. NMR data was analyzed using software MestReNova Lite SE (Mestrelab Research), following manufacturer’s instructions.

### Data availability

The authors declare that the data supporting the findings of this study are available within the paper and its supplementary information files. Any additional information required to interpret the findings of this study is available upon request from the corresponding author (JWL).

